# dialogi: Utilising NLP with chemical and disease similarities to drive the identification of Drug-Induced Liver Injury literature

**DOI:** 10.1101/2022.03.11.483929

**Authors:** Nicholas M Katritsis, Anika Liu, Gehad Youssef, Sanjay Rathee, Méabh MacMahon, Woochang Hwang, Lilly Wollman, Namshik Han

**Author notes:** Correspondence: Milner Therapeutics Institute, Jeffrey Cheah Biomedical Centre, University of Cambridge, Puddicombe Way, Cambridge, CB2 0AW, UK.

## Abstract

Drug-Induced Liver Injury (DILI), despite its low occurrence rate, can cause severe side effects or even lead to death. Thus, it is one of the leading causes for terminating the development of new, and restricting the use of already-circulating, drugs. Moreover, its multifactorial nature, combined with a clinical presentation that often mimics other liver diseases, complicate the identification of DILI-related literature, which remains the main medium for sourcing results from the clinical practice and experimental studies. In this work– contributing to the ‘Literature AI for DILI Challenge’ of the Critical Assessment of Massive Data Analysis (CAMDA) 2021– we present an automated pipeline for distinguishing between DILI-positive and negative papers. We used Natural Language Processing (NLP) to filter out the uninformative parts of a text, and identify and extract mentions of chemicals and diseases. We combined that information with small-molecule and disease embeddings, which are capable of capturing chemical and disease similarities, to improve classification performance. The former are directly sourced from the Chemical Checker (CC). For the latter, we collected data that encode different aspects of disease similarity from the National Library of Medicine’s (NLM) Medical Subject Headings (MeSH) thesaurus and the Comparative Toxicogenomics Database (CTD). Following a similar procedure as the one used in the CC, vector representations for diseases were learnt and evaluated. Two Neural Network (NN) classifiers were developed: one that only accepts texts as input (baseline model) and an augmented classifier that also utilises chemical and disease embeddings (extended model). We trained, validated, and tested the models through a Nested Cross-Validation (NCV) scheme with 10 outer and 5 inner folds. During this, the baseline and extended models performed virtually identically, with macro F_1_-scores of 95.04 ± 0.61% and 94.80 ± 0.41%, respectively. Upon validation on an external, withheld, dataset, representing imbalanced data, the extended model achieved an F_1_-score of 91.14 ± 1.62%, outperforming its baseline counterpart, which got a lower score of 88.30 ± 2.44%. We make further comparisons between the classifiers and discuss future improvements and directions, including utilising chemical and disease embeddings for visualisation and exploratory analysis of the DILI-positive literature.

## 1 INTRODUCTION

Drug-Induced Liver Injury (DILI) is a rare adverse drug reaction that can cause severe complications and, in some cases, may even prove fatal. The term is primarily used to signify the unexpected harm that a drug can cause to the liver. Virtually every class of medication can lead to hepatotoxicity, but the relative risk varies greatly between different drugs (David and Hamilton, 2010). For example, studies suggest that oral medications with doses higher than 50 mg/day and greater lipophilicity– thus those exhibiting higher hepatic metabolism– are more likely to cause DILI (Fontana, 2014; David and Hamilton, 2010).

Liver toxicity can be brought about in a predictable, dose-dependent, manner, when an individual is exposed to concentrations exceeding a drug’s toxicity threshold. This is known as intrinsic (or direct) DILI, has a relatively short latency period (hours to days), and is reproducible in animal models. The most often studied example of intrinsic DILI is acetaminophen (paracetamol), which accounts for about, or more than, half of acute liver failure (ALF) cases in the UK and USA (Katarey and Verma, 2016; Andrade et al., 2019). The majority of DILI cases, however, belong in the idiosyncratic (or indirect) variety, which, as the name suggests, cannot be solely explained by the drug in question. This type of DILI is instead driven by a mixture of characteristics that are unique to the individual and their environment, and tends to have a longer latency period following exposure (days to months) (Andrade et al., 2019). Idiosyncratic DILI is most prominently associated with antibiotics, and amoxicillin-clavulanate is the most commonly implicated drug in studies of European and American populations (Katarey and Verma, 2016).

Idiosyncratic DILI is, indeed, a rare occurrence, with two prospective population-based studies in France (Sgro et al., 2002) and Iceland (Björnsson et al., 2013) placing its crude annual incidence rate at 13.9 and 19.1 cases per 100, 000 people, respectively. A retrospective study of the UK-based General Practice Research Database (GPRD) (de Abajo et al., 2004) reports a lower rate of 2.4 cases per 100, 000 people, which is also in line with other studies from Sweden and the USA (Andrade et al., 2019). Out of those cases, an analysis of data coming from the Spanish DILI registry showed that about 4.2% progress to ALF (Robles-Diaz et al., 2014). This is in agreement with an incident rate of 1.02 cases per 1, 000, 000 people, reported by another US-based study (Goldberg et al., 2015). Yet, despite its rarity, DILI remains one of the commonest reasons for the premature termination of drug development, while also affecting already-circulating drugs, often leading to withdrawal from the market, or issuing warnings and modifications of use (Katarey and Verma, 2016; Andrade et al., 2019). Therefore, the ability to reliably identify cases of DILI in the literature becomes critical, as such resources can aid both physicians in diagnosing the disease and researchers in, among other things, unravelling its mechanisms of action.

The identification of DILI-related literature is complicated by the heterogeneous and multifactorial nature of it. Typically, a drug causes hepatotoxicity directly, through its metabolites, and/or due to possible subsequent inflammatory reaction. However, factors including pre-existing liver pathology, such as Hepatitis B or C, or non-alcoholic fatty liver disease (NAFLD), and chronic alcohol consumption can increase an individual’s susceptibility to DILI. Similarly, genetic factors are at play; different cytochrome p450 enzyme phenotypes can lead to either decreased metabolism of toxic drugs or accelerated production of toxic intermediates, and human leukocyte antigen (HLA) polymorphisms may cause enhanced immune-mediated mechanisms. Furthermore, the clinical presentation of the disease is broad, with symptoms that often mimic other acute and chronic liver diseases, and, in the absence of diagnostic tests and biomarkers, diagnosis is primarily based on establishing a temporal association between drug exposure and symptom development, which is assessed alongside clinical history, liver biochemistry, imaging, and, in some cases, biopsy. (David and Hamilton, 2010; Katarey and Verma, 2016) This complex landscape makes DILI identification a challenging task, with the application of text-mining techniques on DILI-related literature (Cañada et al., 2017; Wu et al., 2021) remaining relatively sparse.

This work presents a contribution to the ‘Literature AI for DILI Challenge’, which was part of the Critical Assessment of Massive Data Analysis (CAMDA) 2021 (http://camda2021.bioinf.jku.at). The aim of the challenge was to develop a classifier capable of identifying DILI-relevant papers. For that, we were given access to about 7, 000 DILI-positive PubMed papers, referenced in the National Institutes of Health’s (NIH) LiverTox database (Hoofnagle et al., 2013), and a non-trivial reference dataset of around 7, 000 DILI-negative papers. These originated from a larger collection of positive and negative corpora that was split in half to create a second dataset, similar in size with the one released (about 14, 000 texts in total), that was withheld and used for final performance testing. We refer to this as ‘external validation’ to distinguish it from the (internal) Nested Cross-Validation (NCV) that we perform. A second, smaller, but more challenging, external validation dataset of 2, 000 papers was also provided.

We built an analysis pipeline that combines Natural Language Processing (NLP) with small-molecule and disease similarities. We pre-processed and normalised the texts to exclude uninformative words and allow for comparisons to be drawn across them. Within each text, chemical and disease terms were annotated and extracted. We treated those as external features and applied a framework that is capable of capturing their similarity. For chemicals, we acquired vectors (embeddings) directly from the Chemical Checker (CC) (Duran-Frigola et al., 2020). For diseases, we first collected data that encode the relations that exist between them. These were sourced from the National Library of Medicine’s (NLM) Medical Subject Headings (MeSH) thesaurus (https://meshb.nlm.nih.gov/) and the Comparative Toxicogenomics Database (CTD) (Davis et al., 2021). We then followed a similar procedure as the one used in the CC to learn vector representations for diseases. Since, typically, a text is associated with multiple terms, an average chemical-and disease-vector (external feature vector) was calculated and attached to it. These, together with the normalised texts, were fed into a Neural Network (NN) classifier. To prevent over-fitting during training, and to get an unbiased estimate of classification performance, we did hyperparameter tuning in a NCV scheme with 10 outer and 5 inner folds. During model evaluation, the extent to which external features alone are capable of distinguishing between the DILI-positive and negative texts was examined. Classifiers with and without the inclusion of external feature vectors were built and compared. During discussion, we explore drawbacks, point out future improvements, and focus on the potential impact of this work on facilitating DILI research.

## 2 METHODS

This analysis is split in three consecutive stages, with each being dependant on the output of the previous ones. First, title and abstract pairs (texts) were collected and processed. This stage constitutes the NLP pipeline, which can be further split in two steps: text pre-processing, and chemical and disease term (concepts) annotation. At the second stage, drug and disease embeddings were learnt, and an average drug- and disease-representation (external feature vector) was calculated for each text. Lastly, the NN classifiers were built, and then trained, validated, and tested in a NCV scheme. The project has been developed in Python 3.9.10 and bundled as a package, to provide ease of use and aid future development.

### 2.1 NLP pipeline

#### 2.1.1 Text pre-processing

Titles and abstracts were first concatenated to form ‘full’ texts. These were then provided as input to the Stanza NLP package (Qi et al., 2020), which was initialised with its tokenisation, lemmatisation, and Part-of-Speech (POS) processors. Stanza provides two biomedical Universal Dependencies (UD) models that are pre-trained on human-annotated treebanks. For this analysis, we used the option that is based on the GENIA corpus (Kim et al., 2003), as it is built on top of 2, 000 PubMed abstracts, and was therefore thought to be a better fit for the (also PubMed-sourced) texts that we had at our disposal.

Each text was split to sentences and then words, and each word was mapped to its base form (lemma). We filtered out lemmas that were not nouns, verbs, adjectives, or adverbs. A list of stopwords was compiled by merging those included in the spaCy package in Python, with the ones provided by PubMed (https://pubmed.ncbi.nlm.nih.gov/help/#help-stopwords). Subsequently, both stopwords and any lemmas that were less than 3 characters long were purged. As a result of those pre-processing steps, implicitly, the texts were also lowercase-normalised, and any numerals and punctuation marks were dropped.

#### 2.1.2 Concept annotation

We queried PubTator Central’s (Wei et al., 2019) RESTful API to acquire annotations for chemicals and diseases. The tool performs concept disambiguation, which resolves conflicts when overlapping annotations are found, and returns concepts normalised to their respective MeSH identifiers. We then counted the times each annotated term shows up within a text, and calculated and assigned (text-specific) relative frequencies to them. In the code, the ‘PubTator’ class is responsible for handling POST and GET requests to the server, processing the raw response data, and associating texts with annotated terms and their relative frequencies. At this step, raw, unchanged, texts were used as input. As a result, the annotations we got back were incompatible, and thus could not be utilised together, with the pre-processed texts of the previous section.

To resolve this issue, while also retaining clarity, the ‘UnivTextBlock’ class was implemented in the code. The class provides a method for exporting processed text, in the sense that pre-processing has been applied and concept terms have been normalised, either by replacement with their MeSH identifiers, or the broader concept category (that is, ‘disease’ or ‘chemical’). To handle cases where a concept term spans across multiple words, or is misaligned compared to the target word(s)– often the result of incorrect sentence segmentation or peculiarities in tokenisation– the code checks for degree of overlap. We observed good performance when demanding that the latter exceed a minimum threshold value of 90%.

### 2.2 External feature vector generation

#### 2.2.1 Data collection

We aimed to quantify disease and, separately, chemical similarity. First, we collected data from the MeSH thesaurus. Descriptors and supplementary concept records were downloaded in XML format. At the uppermost level, there are 16 categories, which are further split in subcategories. Within each subcategory, descriptors are arranged in a hierarchical manner, from most general to most specific. This results in a branching, tree-like, structure. In the XML file, each descriptor is associated with one or multiple tree numbers, which represent paths taken from the root subcategory, until the descriptor in question is reached.

These data were parsed into a dictionary that associates descriptors with their respective tree numbers. We selected for disease-descriptors by pruning those, whose trees did not start with ‘C’, as category C is for diseases. Similarly, for chemicals, we filtered out descriptors that did not fall under category D, which contains drugs and chemicals. Supplementary concept records are not associated with tree numbers. Instead, they are mapped onto one or multiple descriptors. We parsed these relationships in a separate dictionary, which we used to indirectly link supplementary concepts with the hierarchical structure described earlier.

We also collected data from the CTD, which associates diseases with chemicals, genes, pathways, and phenotypes. In this context, phenotypes refer to non-disease biological events and are expressed using the Gene Ontology (GO) as controlled vocabulary (Davis et al., 2021). As a result, disease-phenotype associations are further split into three datasets, one for each GO category: Biological Process (GO-BP), Cellular Component (GO-CC), and Molecular Function (GO-MF). In total, 6 datasets were downloaded from the CTD. Disease terms are expressed using MeSH identifiers and, thus, required no further processing.

#### 2.2.2 Concept embedding learning

After data collection, a procedure matching the one followed in the CC was used. This was applied separately to each dataset and consists of three consecutive steps: (a) turning the dataset into a corpus, (b) learning a (sparse) vector representation on it, and (c) embedding the latter into a lower dimensional space. For the MeSH, concept terms are associated with tree numbers, which represent paths. We traversed these paths, starting from the root subcategory, and saved each location on the tree as a word. A corpus was, then, built by repeating this process for all terms. For the CTD, concept terms are linked together through interactions with chemicals, genes, etc. We created a corpus by treating these interacting partners as words.

Corpora were mapped to a sparse vector space by applying a term frequency–inverse document frequency (TF-IDF) transformation. Prior to that, frequent and infrequent words were dropped, that is, words associated with less than 5 or more than 80% of the terms. Following the transformation, any null (zero) vectors were purged, alongside with their corresponding terms. An initial dimensionality reduction step was performed by means of truncated Singular Value Decomposition (SVD). Here, we kept the number of components that explained around 90% of the variance seen in the original data.

For learning the final embeddings, we ran the node2vec algorithm (Grover and Leskovec, 2016), with its default parameters, on a term similarity network. This produced 128-dimensional vectors. To create the similarity network, we first used cosine similarities to identify each term’s neighbourhood, which consisted of its 100 closest neighbours. We approximated the null distribution empirically, by randomly sampling with replacement 100, 000 pairs of terms and calculating their cosine similarities. This was used to map neighbour similarities to p-values. We built the network by merging the neighbourhoods together and assigning *−log*_10_(p-values) as edge weights. We pruned any insignificant edges– that is, edges with weights less than, or equal to, 2– but demanded that each term be connected to at least 3 closest neighbours.

#### 2.2.3 Chemical Checker embeddings

While chemical similarities from the CC need no further processing themselves, vectors are indexed by their InChIKeys. Since we normalise chemicals using MeSH identifiers, a mapping had to be created that would link the two. We queried ChemIDplus’s (https://chem.nlm.nih.gov/chemidplus/) API to retrieve MeSH terms and their respective InChIKeys and SMILES. We, then, used the MeSH thesaurus to associate MeSH identifiers with concepts and terms. However, a MeSH identifier usually points to multiple concepts– which typically consist of one or more synonymous terms– and sometimes more than one of those concepts or terms are associated with an InChIKey and/or SMILES. Therefore, to reliably translate from MeSH to InChIKeys, we also took into account the hierarchy of preferred concepts and terms that exists in the MeSH thesaurus.

In an attempt to further expand the number of MeSH terms that are associated with an InChIKey, we extracted relevant associations from the DrugBank database (Wishart et al., 2018) and merged them with those associations sourced from ChemIDplus. Furthermore, in order to enrich any sparse CC spaces, we utilised the CC signaturisers (Bertoni et al., 2021) to predict embeddings for chemical compounds that are not included in the original database. Here, the SMILES structural information– which had been acquired from ChemIDplus in the previous step– were given as input to the signaturisers.

#### 2.2.4 External feature vectors

##### 2.2.4.1 Generation

At this stage, disease embeddings have been generated– seven vector spaces in total; six from the CTD and another one from the MeSH– and chemical signatures have been retrieved from the Chemical Checker (25 spaces, augmented with an additional one, generated from MeSH data). We first aimed to concatenate individual, concept-specific, spaces into one, so that diseases (and chemicals) were represented by a single vector space (and chemicals by a separate one). However, concatenation alone would not only lead to a considerable dimensionality difference between the resulting disease- and chemical-specific spaces (896 and 3,328 dimensions, respectively), but also potentially combine a great number of correlated features together, adding redundant dimensions to the produced space. Therefore, after concatenation, followed a dimensionality reduction step using truncated SVD. We have carefully tuned this process to retain as much information as possible, without introducing additional noise.

The vectors were not normalised or centred prior to concatenation and dimensionality reduction, as doing so, in this particular case, did not lead to significant differences. For diseases, we chose to concatenate the top two most dense CTD spaces (GO-BP and GO-MF, for rationale, see Results) and then performed truncated SVD to reduce them down to 103 dimensions, which explained about 90% of the original variance. Separately, we reduced the dimension of the MeSH space for diseases from 128 to 47 dimensions, which also retained about 90% of the variance of the original data. We concatenated the two reduced spaces to form the final 150*−*dimensional disease space. In a similar manner, for chemicals, we concatenated all the CC spaces and reduced them down to 265 dimensions. The MeSH space for chemicals was also reduced to 35 dimensions. In both cases, about 80% of the original variance was explained. By concatenation, the final chemical space was produced, which is 300*−*dimensional.

At this point, a single disease-specific, and a second chemical-specific, space exists. These are by no means related to the texts, but instead encode similarities between the concepts that were extracted out of them earlier. In contrast, external feature vectors are meant to be text-specific and to capture the similarities between the texts, as these are encoded by the combinations of chemicals and diseases that show up in them. For each text, concept relative frequencies– already calculated during concept annotation– were used to calculate the weighted average for the disease embeddings and, separately, chemical embeddings. Concept embeddings that belong to terms mentioned infrequently within the text are, as a result, down-weighted, compared to those associated with more frequent terms. The two (now text-and concept-specific) vectors were first normalised to unity and then concatenated to form the final external feature vector.

##### 2.2.4.2 Comparisons

For between-space comparisons, two measures were used: the Rank-Biased Overlap (RBO) (Webber et al., 2010), and Pearson correlation. The RBO is a top-weighted similarity measure, that can be applied to non-conjoint ranked lists of indefinite length. The measure models the behaviour of a user comparing between two lists incrementally, at increasing depths, where, at each depth, a fixed probability of stopping exists.

To compare the similarity between two spaces, a procedure similar to the one described in the Chemical Checker was followed (Duran-Frigola et al., 2020). First, the common concepts between the two spaces were identified. Then, for each concept, we computed a list of its 100 closest neighbours. We used cosine similarities and returned lists that were ordered by decreasing similarity. The similarity search was performed efficiently using the Faiss library in Python (Johnson et al., 2017). The two ranked lists were used to calculate a RBO similarity score (we set *p* = 0.7, making the search more top-weighted). The process was repeated for all common concepts, the similarity scores were aggregated, and their average value was calculated. This was used as the RBO similarity score for the space-pair.

To calculate Pearson correlations, we applied Canonical Correlation Analysis (CCA) on space-pairs to find orthogonal linear combinations (canonical variables) of their features, that maximally correlate with each other. We kept the first three canonical variable pairs and, for each of them, calculated the Pearson correlation. By averaging those values together, the final space-pair correlation value was calculated.

### 2.3 NN classifiers and validation

We developed two NN classifiers. The baseline classifier accepts processed texts as its single input. The extended classifier augments the baseline model by additionally taking into account each text’s external feature vector (Fig. 1). First, the texts are fed into an embedding layer, which has been initialised with GloVe vectors (Pennington et al., 2014). Then, these embeddings pass through a Bidirectional Long Short-Term Memory (BiLSTM) layer, which is followed by a dense layer with ReLU activation. An output dense layer with sigmoid activation is used to compute the classification probability value.

**Figure 1.**
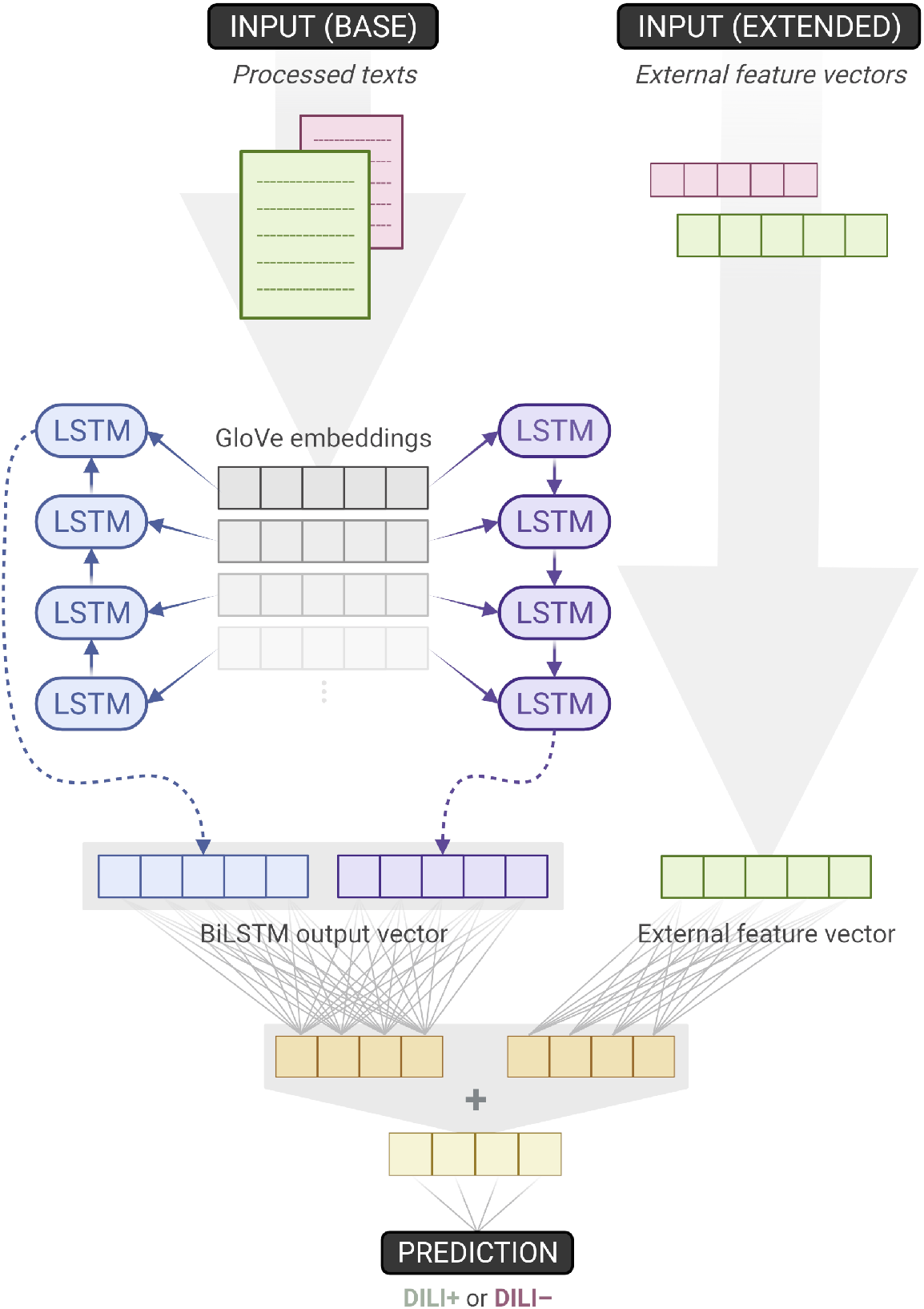
Overview of the baseline and extended classifiers. The former accepts texts as its single input, while the latter augments its baseline counterpart by also utilising external feature vectors. When training the baseline model, weights downstream the ‘base input’– on the left side of the figure– are learnt. As the extended model is built on top of the baseline, those weights can be transferred and frozen, thus remaining unchanged during training. As a result, for the extended model, only one dense layer’s weights have to be trained. (Created with BioRender.com)

For the extended classifier, the external feature vectors first pass through a separate ReLU dense layer that is chosen to have the same number of units as the one mentioned earlier. Thus, the outputs of those two dense layers can be added together, before going through the same sigmoid output dense layer, as the one used in the baseline model. This design choice is intentional, it introduces no additional hyper-parameters to optimise, and allows for both the baseline and extended models to be trained and tested within the same NCV scheme. We trained the 10 outer-fold baseline models (which were inner-fold winners), froze their weights, augmented and transformed them to extended models, and then repeated the training one more time. As a result, for the extended model, just a single dense layer’s weights needed to be trained.

During initial testing and tuning, it became apparent that using a Bi-instead of a Uni-LSTM layer, and allowing for the text embeddings to be trainable, consistently led to better performing models. Therefore, we did not optimise for those parameters. Nonetheless, hyper-parameter tuning was applied within a NCV scheme with 10 outer and 5 inner folds. We varied the embedding dimension ([50, 100, 200, 300]), UniLSTM units (32 *−* 96, with a step size of 16), dense layer units (192 *−* 320, with a step size of 32), and the learning rate ([10^−3^, 5 *×* 10^−3^, 7 *×* 10^−3^, 10^−2^]). During model training, we used a batch size of 32, and the Adam optimiser with binary cross-entropy as the loss function. For hyper-parameter tuning, we monitored validation loss. When training the extended model, a fixed learning rate of 10^−2^ was used.

Additionally, we observed that the models learn rapidly, and usually start to overfit within the first 10 epochs, even with dropout and L1/L2 regularisation applied appropriately to the LSTM and dense layers. In fact, training for just one epoch tended to produce models performing similarly, or better, than those trained for longer. Thus, we chose to train for no more than one epoch. In this case, regularisation does not improve performance, and is thus omitted (Komatsuzaki, 2019). We use Keras and the Bayesian optimisation algorithm in KerasTuner to build, train, and validate the NN classifiers, and to perform hyper-parameter optimisation. To support NCV, the original KerasTuner code was subclassed and extended. We generate stratified k-folds through Scikit-learn’s (Pedregosa et al., 2012) ‘StratifiedKFold’ function.

## 3 RESULTS

### 3.1 Concept embeddings

We first examined the degree of term coverage for the different concept spaces (Fig. 2A). Chemical and disease terms found in the texts have already been extracted and collected. However, there are terms missing in some of the spaces. This is either due to the term not being present in the data that was used to construct the spaces in the first place, or a result of the TF-IDF word filtering steps that were applied afterwards.

**Figure 2.**
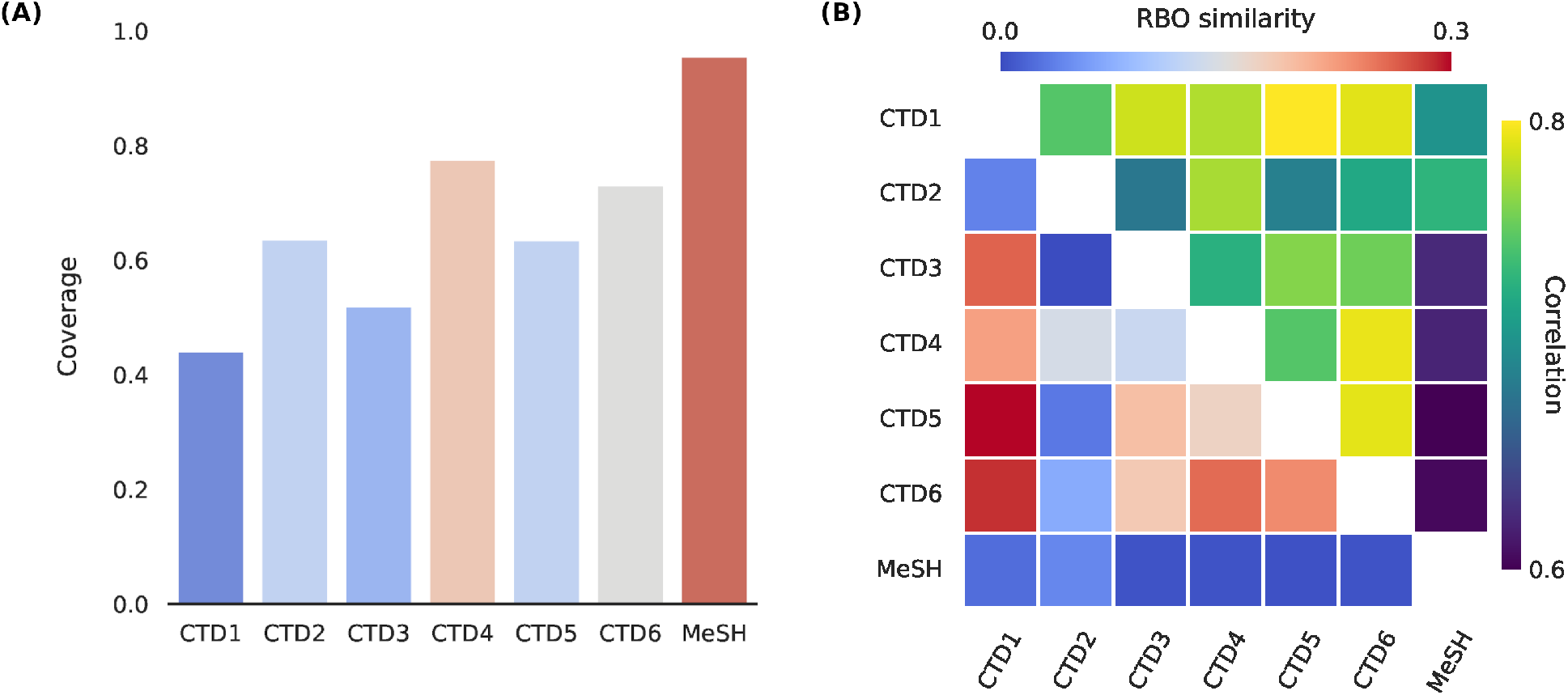
CTD1-6: Genes, Chemicals, Pathways, GO-BP (Biological Process), GO-CC (Cellular Component), and GO-MF (Molecular Function). **(A)** Disease term coverage for the 6 CTD (Comparative Toxicogenomics Database) spaces, and MeSH. The latter, together with GO-BP and GO-MF, are the most enriched spaces, encoding about 96%, 78%, and 73%, respectively, of all the disease terms found in text. **(B)** RBO (Rank-Biased Overlap) similarities and Pearson correlations between the different disease spaces. Although the two are not directly comparable, they seem to be in good agreement with each other, with MeSH being the least, and CTD1 (Genes) the highest correlated space, in general.

Missing terms are represented as null vectors. For diseases, MeSH is the most enriched space (96% coverage), followed by the CTD’s GO-BP and GO-MF (coverage of 78% and 73%, respectively). Since we normalise terms to the MeSH vocabulary, the MeSH space is expected to be the most enriched.

For chemicals, the CC spaces appear to be equally enriched, with a coverage of about 60%. The MeSH space for diseases has a higher coverage of 84%. The uniformity that is observed across the different CC spaces can be attributed to, and also supports, the usage of the CC signaturisers. These fill in the gaps of missing molecular signatures; typically, CC spaces tend to differ considerably in terms of their sizes (Duran-Frigola et al., 2020). Notably, a concept space with lower term coverage does not necessarily translate to external feature vectors with reduced text coverage. The latter are (text-specific) linear combinations of concept vectors, and the coverage of that space is, thus, also affected by the combination of terms that show up in each particular text, as well as their relative frequencies.

We then calculated the RBO similarity measures and Pearson correlations across the different pairs of disease spaces (Fig. 2B). The two measures are in good agreement with each other. As expected, given that the rest of the spaces are based on the CTD-sourced datasets, the MeSH space tends to be the most dissimilar one, followed by the CTD’s Chemicals space. On the other end, CTD Genes is highly correlated with most other CTD spaces. We created similar plots to compare between the chemical spaces and, for the CC spaces, observed a similarity and correlation profile that matched the one provided and discussed in the original publication (Duran-Frigola et al., 2020).

We utilised both coverages and correlations when selecting for the disease spaces and, separately, chemical spaces, to concatenate. For diseases, we chose the top three enriched spaces (MeSH, and CTD GO-BP and GO-MF). When seen as a group, these are strongly correlated with the rest of the CTD spaces. We chose to concatenate all chemical spaces together. The premise here is that concatenation between spaces with largely different term coverages might introduce unwanted noise later, during the dimensionality reduction step (see Methods). This is of concern for diseases, where coverage ranges from as low as 44% to a highest score of 96%, but not for chemicals, where it remains virtually unchanged across the CC spaces.

### 3.2 External feature vectors

We were also interested in assessing the extent to which the external feature vectors are capable of capturing the differences between DILI positive and negative texts. Ability to do so, at this stage, would provide strong evidence of their suitability to be used as additional inputs to the (extended) classifier. First, we normalised the vectors, performed Principal Component Analysis (PCA), and kept the first 15 components. We, then, produced a 3D t-SNE plot (Fig. 3). Throughout this work, as is usual in NLP, we are working with cosine similarities. However, cosine distances are not invariant to mean-centring– which PCA explicitly performs– and will be affected and distorted. In contrast, euclidean distances are mean-centring invariant. By normalising the data first, we enforce a monotonic relationship between cosine and euclidean distances, which we exploit by using euclidean distances in the t-SNE plot (Korenius et al., 2007).

**Figure 3.**
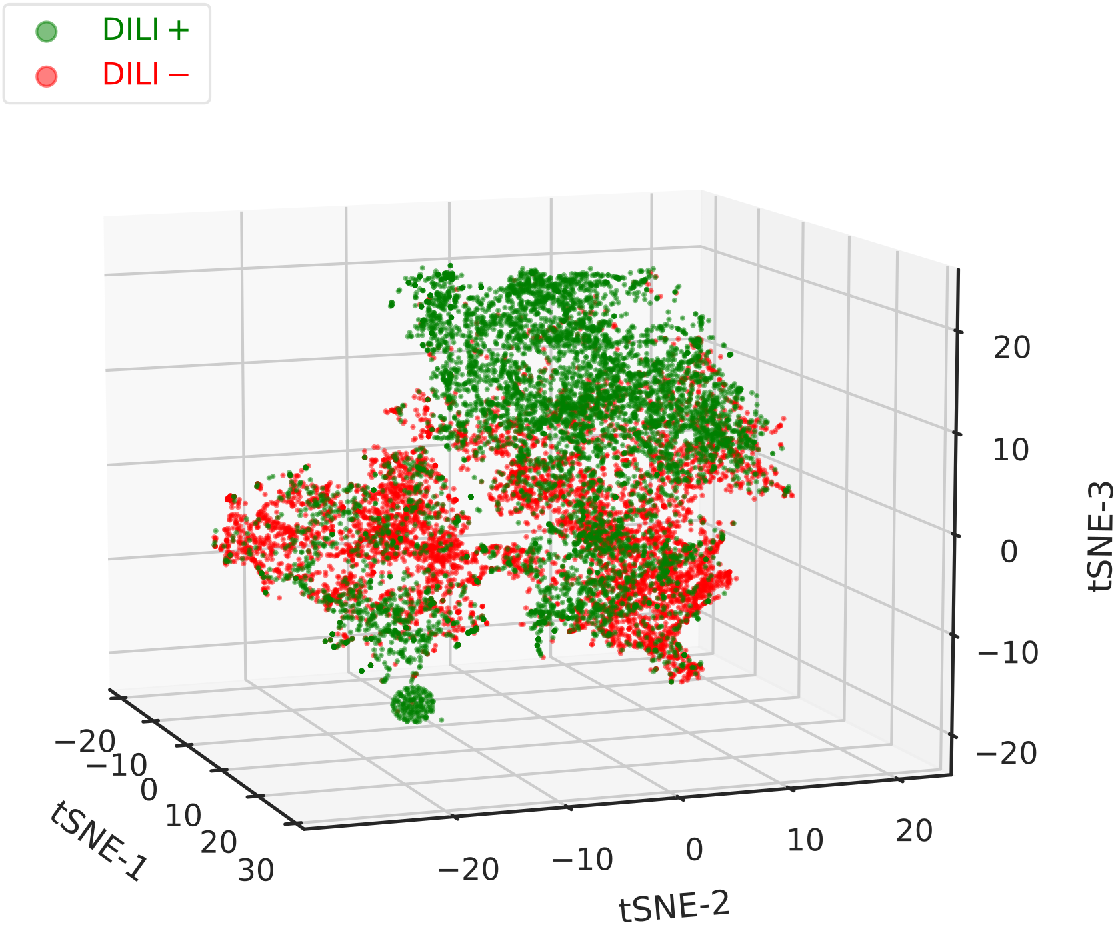
3D t-SNE plot of all the non-zero external feature vectors (which combine both chemical and disease embeddings). A good degree of separation can be observed between the DILI positive and negative texts, with the former clustering cleanly on the top-right corner of the plot. A second dense cluster is formed at the bottom-left corner. Between them, in the middle part of the plot, reside both positive and negative texts, that did not form distinct clusters. While feature vectors can improve classification performance, they tend to be sparse, and should, therefore, not used as the single input to the classifier.

In the plot (Fig. 3), a good degree of separation can be observed between the DILI positive and negative samples. Positive texts tend to cluster in the upper-right, and also form a tight cluster in the lower-left, corner. Between those, both positive and negative texts reside, forming largely overlapping clusters. It should be pointed out that concept vectors were learnt completely separately from, and are in no way connected with, the classification of texts in the two classes. As such, the acceptable clustering performance seen here should be attributed in: (a) similar chemicals and/or diseases appearing within each class, (b) dissimilar chemicals and/or diseases appearing between classes, (c) unique chemical, disease, and chemical-disease combinations dominating in each class. This might be worth further investigation, but, for now, makes an appealing case for the usefulness of the external feature vectors as a means for improving classification performance.

### 3.3 Classification performance

We compared between the baseline classifier, which uses texts as its sole input, and the extended one, that also accepts external feature vectors. As we are interested in the balance between precision and recall, we used the F_1_-score as performance measure. During internal validation, macro F_1_-score (average of per-class scores) was calculated. For external validation, micro scores (calculated over the entirety of the predictions, irrespective of classes) are reported. During initial tuning and testing, we observed that the baseline model performs optimally with the usual classification threshold of 0.5, but, for the extended model, a higher threshold of about 0.7 leads to unchanged, or improved, performance, depending on the validation dataset used. We set these, seemingly arbitrary, thresholds at the beginning of the NCV procedure and evaluated their suitability afterwards. Alternatively, a more elegant approach would treat the classification threshold as a hyper-parameter to be optimised in the inner NCV folds.

We plotted the average performance across the 10 outer folds (Fig. 4). During internal validation, the baseline and extended models performed virtually identically, with F_1_-scores of 95.04 ± 0.61% and 94.80 ± 0.41%, respectively. Evaluating the models on the first external dataset, which represents balanced data, painted a similar picture; this also provides proof that the training procedure we utilise does not lead to overfitting. In this case, baseline and extended models achieved scores of 95.11 ± 0.34% and 94.93 ± 0.48%, respectively. We observed a drop in performance, which affected both models, when testing on the second external dataset, that represents imbalanced data. However, the extended model managed to outperform the baseline model by a considerable margin; the former achieved an F_1_-score of 91.14 ± 1.62%, compared to the baseline’s 88.30 ± 2.44%. The extended classifier also seems to produce lower dispersed scores, which becomes especially pronounced during the second phase of external validation.

**Figure 4.**
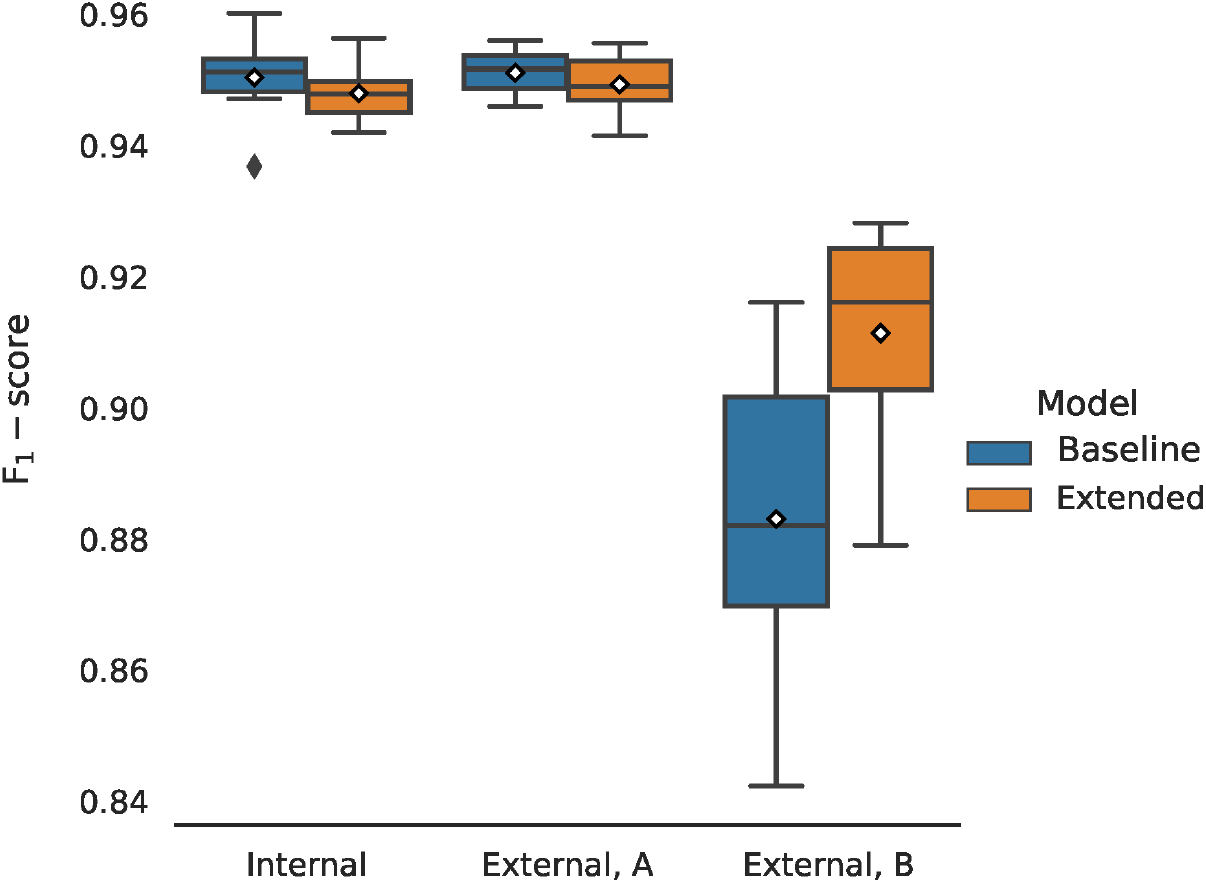
Classification performance comparison between the two models, for different validation datasets. The performance across 10 folds is being reported. Mean values are annotated with white diamonds. Both models perform virtually identically during internal validation (macro F_1_-scores of 95.04 ± 0.61% and 94.80 ± 0.41%, respectively), and equally well when tested on the first external dataset (micro F_1_-score of 95.11 ± 0.34% and 94.93 ± 0.48%, respectively). The models are not overfitting. There is a significant drop in performance when testing on the second external dataset (which simulates imbalanced data). However, the extended model performs considerably better, with a (micro) F_1_-score of 91.14 ± 1.62%, compared to the baseline’s 88.30 ± 2.44%.

Lastly, we evaluated the choice of threshold for the two classifiers. Within each outer fold, we varied the threshold between 0.5 and 0.95 and calculated the (macro) F_1_-scores (Fig. 5). When compared at the same classification threshold, the extended model consistently outperforms its baseline counterpart by a small margin, at thresholds closer to 0.5, which incrementally grows larger at higher thresholds. This implies a difference between the slopes of the two curves which is, indeed, there to be seen: the baseline curve is steeper at each threshold value, compared to the extended one. The inclusion of the external feature vectors has resulted in the extended classifier being more confident in its predictions, which is reflected in the probability scores being pushed closer to the limit points of the [0, 1] interval (and a lower binary cross-entropy validation loss, too). It is desirable to set the threshold to a higher value, as doing so can improve– sometimes considerably– the classification performance on the imbalanced external dataset. Higher thresholds, however, might hurt the performance on the balanced validation datasets. For the extended model, choosing a threshold in the range of 0.5 *−* 0.7 leads to a virtually unchanged F_1_-scores, a behaviour not followed by the baseline model. With this in mind, the choice of thresholds for the two classifiers seems to be near-optimal.

**Figure 5.**
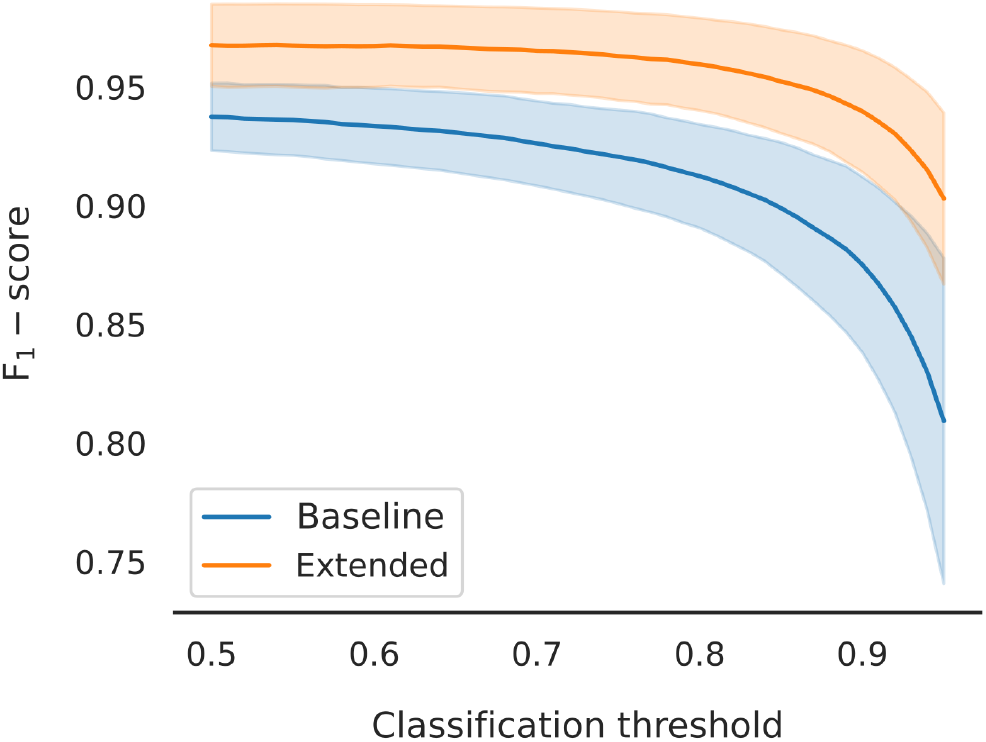
Macro F_1_-score as a function of classification threshold, varied in the domain of [0.5, 0.95]. Mean values and standard deviations are plotted. The extended model outperforms the baseline at every threshold value. There is a clear difference in slope, with the baseline curve being steeper throughout the domain. Learning from the external feature vectors has pushed the extended model to be more confident in its predictions. In turn, using the same threshold for both models will lead to (at least) one of them under-performing. A higher threshold tends to improve the performance of the models on imbalanced data.

## 4 DISCUSSION

Word embeddings learnt directly on DILI-related (or other biomedical) literature cannot capture the similarities between diseases and chemicals. GloVe embeddings, for example, encode linguistic and/or semantic similarities of words by taking into account co-occurrences. However, there are no semantics to be encoded when it comes to chemical names or diseases in regular text. Of course, chemical (and disease) terms can still be similar, and this similarity could still be expressed in terms of co-occurrences, but instead of words in regular text, one would use common protein targets, gene pathways, indications, etc., to acquire meaningful relations. This is the rationale behind the usage of concept embeddings in this work.

Concept embeddings, turned into text-specific external feature vectors, however, present a challenge when utilised alone for classification. For the DILI-positive class, 93% and 86% of the texts have been annotated with at least one disease and chemical term, respectively. In the negative class, these percentages shrink down to 78%, for diseases, and 56%, for chemicals. The lack of annotated concept terms can be attributed to: (a) failure to annotate terms that exist in the text (false negatives), or (b) genuine lack of terms (true negatives), or (c) lack of terms in the title and/or abstract, but presence in the full text (true negatives in the context of the challenge, but false negatives in the broader sense). Because of the first and last points, filtering out texts with no annotated concepts as DILI-negative would be problematic. Instead, combining concept with word embeddings enables the classifier to make informed decisions, even when no chemicals or disease terms have been identified. Then, acquiring full texts when no concept terms are included in the title and abstract, could have the potential to improve classification performance.

The additional information that we either generate, or collect, about disease, chemical, and text similarity can also prove valuable for the purposes of visualisation and exploratory analysis. Similar to the t-SNE plot that we provide in this study (Fig. 3), texts could be further clustered together based on the combination of chemicals or, alternatively, diseases that occur therein– a process that inherently takes into account concept similarities, too. Alternatively, average chemical similarities could be calculated against drugs that are already known to cause DILI, for example with the help of the DILIrank dataset (Chen et al., 2016). These could then be used to rank DILI-positive texts from most (high similarity to known DILI-related drugs) to less promising, as well as annotate them separately on the t-SNE plot, so that their neighbourhoods can be identified and further explored. This is one of the most exciting future prospects of this work.

Overall, in this study, we have demonstrated that utilising disease and chemical embeddings, alongside a typical NLP pipeline, has the potential to considerably improve classification performance. On external validation, the best performing classifier achieved an average F_1_-score of 94.93 ± 0.48%, on balanced data, and of 91.14 ± 1.62%, on the dataset representing the imbalanced case. The classifiers’ performance on the latter was tuned with threshold moving and evaluated to showcase that the inclusion of the external features leads to improved and more consistent performance. We also demonstrated the capability of the concept embeddings alone to distinguish between the positive and negative literature and discussed their potential usefulness for visualisation and exploratory purposes.

## AUTHOR CONTRIBUTIONS

N.M.K. conceived and designed the study, performed the analysis, collected data, and wrote the manuscript. A.L., G.Y., S.R., M.M., W.H., and L.W. contributed expertise and analysis tools. N.H. supervised the project.

## DATA AVAILABILITY STATEMENT

The code and datasets (directly uploaded or with links to the original sources) used in this study can be found on the GitHub repository: https://github.com/pokedthefrog/camda2021-dialogi

## REFERENCES

Andrade, R. J., Chalasani, N., Björnsson, E. S., Suzuki, A., Kullak-Ublick, G. A., Watkins, P. B., et al. (2019). Drug-induced liver injury. Nat Rev Dis Primers 5, 58. doi:10.1038/s41572-019-0105-0

Bertoni, M., Duran-Frigola, M., Badia-I-Mompel, P., Pauls, E., Orozco-Ruiz, M., Guitart-Pla, O., et al. (2021). Bioactivity descriptors for uncharacterized chemical compounds. Nat. Commun. 12, 3932. doi:10.1038/s41467-021-24150-4

Björnsson, E. S., Bergmann, O. M., Björnsson, H. K., Kvaran, R. B., and Olafsson, S. (2013). Incidence, presentation, and outcomes in patients with drug-induced liver injury in the general population of iceland. Gastroenterology 144, 1419–25, 1425.e1–3; quiz e19–20. doi:10.1053/j.gastro.2013.02.006

Cañada, A., Capella-Gutierrez, S., Rabal, O., Oyarzabal, J., Valencia, A., and Krallinger, M. (2017). LimTox: a web tool for applied text mining of adverse event and toxicity associations of compounds, drugs and genes. Nucleic Acids Res. 45, W484–W489. doi:10.1093/nar/gkx462

Chen, M., Suzuki, A., Thakkar, S., Yu, K., Hu, C., and Tong, W. (2016). DILIrank: the largest reference drug list ranked by the risk for developing drug-induced liver injury in humans. Drug Discov. Today 21, 648–653. doi:10.1016/j.drudis.2016.02.015

David, S. and Hamilton, J. P. (2010). Drug-induced liver injury. US Gastroenterol. Hepatol. Rev. 6, 73–80

Davis, A. P., Grondin, C. J., Johnson, R. J., Sciaky, D., Wiegers, J., Wiegers, T. C., et al. (2021). Comparative toxicogenomics database (CTD): update 2021. Nucleic Acids Res. 49, D1138–D1143. doi:10.1093/nar/gkaa891

de Abajo, F. J., Montero, D., Madurga, M., and García Rodríguez, L. A. (2004). Acute and clinically relevant drug-induced liver injury: a population based case-control study. Br. J. Clin. Pharmacol. 58, 71–80. doi:10.1111/j.1365-2125.2004.02133.x

Duran-Frigola, M., Pauls, E., Guitart-Pla, O., Bertoni, M., Alcalde, V., Amat, D., et al. (2020). Extending the small-molecule similarity principle to all levels of biology with the chemical checker. Nat. Biotechnol. 38, 1087–1096. doi:10.1038/s41587-020-0502-7

Fontana, R. J. (2014). Pathogenesis of idiosyncratic drug-induced liver injury and clinical perspectives. Gastroenterology 146, 914–928. doi:10.1053/j.gastro.2013.12.032

Goldberg, D. S., Forde, K. A., Carbonari, D. M., Lewis, J. D., Leidl, K. B. F., Reddy, K. R., et al. (2015). Population-representative incidence of drug-induced acute liver failure based on an analysis of an integrated health care system. Gastroenterology 148, 1353–61.e3. doi:10.1053/j.gastro.2015.02.050

Grover, A. and Leskovec, J. (2016). node2vec: Scalable feature learning for networks

Hoofnagle, J. H., Serrano, J., Knoben, J. E., and Navarro, V. J. (2013). LiverTox: a website on drug-induced liver injury. Hepatology 57, 873–874. doi:10.1002/hep.26175

Johnson, J., Douze, M., and Jégou, H. (2017). Billion-scale similarity search with GPUs

Katarey, D. and Verma, S. (2016). Drug-induced liver injury. Clin. Med. 16, s104–s109. doi:10.7861/clinmedicine.16-6-s104

Kim, J.-D., Ohta, T., Tateisi, Y., and Tsujii, J. (2003). GENIA corpus–semantically annotated corpus for bio-textmining. Bioinformatics 19 Suppl 1, i180–2. doi:10.1093/bioinformatics/btg1023

Komatsuzaki, A. (2019). One epoch is all you need

Korenius, T., Laurikkala, J., and Juhola, M. (2007). On principal component analysis, cosine and euclidean measures in information retrieval. Inf. Sci. 177, 4893–4905. doi:10.1016/j.ins.2007.05.027

Pedregosa, F., Varoquaux, G., Gramfort, A., Michel, V., Thirion, B., Grisel, O., et al. (2012). Scikit-learn: Machine learning in python

Pennington, J., Socher, R., and Manning, C. (2014). GloVe: Global vectors for word representation. In Proceedings of the 2014 Conference on Empirical Methods in Natural Language Processing (EMNLP) (Doha, Qatar: Association for Computational Linguistics), 1532–1543. doi:10.3115/v1/D14-1162

Qi, P., Zhang, Y., Zhang, Y., Bolton, J., and Manning, C. D. (2020). Stanza: A python natural language processing toolkit for many human languages

Robles-Diaz, M., Lucena, M. I., Kaplowitz, N., Stephens, C., Medina-Cáliz, I., González-Jimenez, A., et al. (2014). Use of hy’s law and a new composite algorithm to predict acute liver failure in patients with drug-induced liver injury. Gastroenterology 147, 109–118.e5. doi:10.1053/j.gastro.2014.03.050

Sgro, C., Clinard, F., Ouazir, K., Chanay, H., Allard, C., Guilleminet, C., et al. (2002). Incidence of drug-induced hepatic injuries: a french population-based study. Hepatology 36, 451–455. doi:10.1053/jhep.2002.34857

Webber, W., Moffat, A., and Zobel, J. (2010). A similarity measure for indefinite rankings. ACM Trans. Inf. Syst. Secur. 28, 1–38. doi:10.1145/1852102.1852106

Wei, C.-H., Allot, A., Leaman, R., and Lu, Z. (2019). PubTator central: automated concept annotation for biomedical full text articles. Nucleic Acids Res. 47, W587–W593. doi:10.1093/nar/gkz389

Wishart, D. S., Feunang, Y. D., Guo, A. C., Lo, E. J., Marcu, A., Grant, J. R., et al. (2018). DrugBank 5.0: a major update to the DrugBank database for 2018. Nucleic Acids Res. 46, D1074–D1082. doi:10.1093/nar/gkx1037

Wu, Y., Liu, Z., Wu, L., Chen, M., and Tong, W. (2021). BERT-Based natural language processing of drug labeling documents: A case study for classifying Drug-Induced liver injury risk. Front Artif Intell 4, 729834. doi:10.3389/frai.2021.729834

